# A theory of direction selectivity for Macaque primary visual cortex

**DOI:** 10.1101/2021.05.02.442297

**Authors:** Logan Chariker, Robert Shapley, Michael Hawken, Lai-Sang Young

## Abstract

This paper offers a new theory for the origin of <u>direction selectivity</u> in the Macaque primary visual cortex, V1. Direction selectivity (DS) is essential for the perception of motion and control of pursuit eye movements. In the Macaque visual pathway, DS neurons first appear in V1, in the Simple cell population of the Magnocellular input layer 4Cα. The LGN cells that project to these cortical neurons, however, are not direction-selective. We hypothesize that DS is initiated in feedforward LGN input, in the summed responses of LGN cells afferent to a cortical cell, and it is achieved through the interplay of (a) different visual response dynamics of ON and OFF LGN cells, and (b) the wiring of ON and OFF LGN neurons to cortex. We identify specific temporal differences in the ON/OFF pathways that together with (b) produce distinct response time-courses in separated subregions; analysis and simulations confirm the efficacy of the mechanisms proposed. To constrain the theory, we present data on Simple cells in layer 4Cα in response to drifting gratings. About half of the cells were found to have high DS, and the DS was broad-band in spatial and temporal frequency (SF and TF). The proposed theory includes a complete analysis of how stimulus features such as SF and TF interact with ON/OFF dynamics and LGN-to-cortex wiring to determine the preferred direction and magnitude of DS.

**Significance Statement:** Motion perception is important for primates, and direction selectivity (DS), the ability to perceive the direction a target is moving, is an essential part of motion perception. Yet no satisfactory mechanistic explanation has been proposed for the origin of DS in primate visual cortex up until now. In this paper, we hypothesize that DS is initiated in feedforward LGN input as a result of the dynamic differences between the ON/OFF pathways. The mechanisms we propose are biology-based, and our theory explains experimental data for all spatial and temporal frequencies in visual stimuli. Exploiting temporal biases in parallel pathways is relevant beyond visual neuroscience; similar ideas likely apply to other types of neural signal processing.g

## Introduction

This paper proposes a solution to a longstanding question in visual neuroscience, namely the origin of direction selectivity in the visual cortex of Macaque monkeys. Motion perception is a vital visual capability well developed in primates. As perceiving motion requires perceiving the direction in which a target moves, direction selectivity (DS), the ability of visual neurons to sense the direction of movement, is essential for motion perception (1) and for the control of pursuit eye movements (2). For these reasons, understanding DS is an important first step towards understanding how the cortex processes motion signals.

Direction selectivity in cortical neurons was first documented in the cat (3). Since then, it has been found in neurons all along the visual dorsal stream (an area associated with motion processing) in primates like Macaque monkeys (4-7), whose vision is like that of humans. DS neurons are in fact present across species; they are widespread among visual mammals, an experimental fact that testifies to their biological significance.

In the visual pathway of Macaques, DS appears first in the primary visual cortex (V1), in the Simple cell population of the input layer 4Cα (8) These neurons provide feedforward DS signals to subsequent cortical layers and brain regions in the dorsal pathway. Thus to discover the origin of DS, one is led to examining how neurons in layer 4Cα acquire their DS ---and that is where it gets interesting: The neurons that provide visual signals to layer 4Cα, the Magnocellular cells in the Lateral Geniculate Nucleus (LGN), are not direction-selective (9-12). Yet many of the cells in the input layer of V1 to which they project are DS. A fundamental scientific question, therefore, is how 4Cα neurons acquire their DS. That is the question we would like to answer in this paper.

Though many papers have been written on DS since its discovery over half a century ago, and there is continued interest in the subject (13-16), no satisfactory mechanistic explanation for the origin of DS in primate cortex has been proposed before now. Early conceptual models of how DS may arise, such as the Reichardt multiplier (17) or the motion-energy model (18), were not concerned with biological mechanisms. Later work proposed neural mechanisms for the motion-energy model (19) but they are not sufficient for explaining DS in primate cortex. See **Discussion** for our comparisons of different model mechanisms.

It is widely accepted that DS computation requires spatio-temporal inseparability or STI, that is, different subregions of the receptive field have different time-courses of response (18, 20, 21). What were lacking were biological mechanisms that could produce STI, and a clear understanding of how DS depends on the interaction between STI and the spatial and temporal character of the visual stimulus. These are the issues we address in this paper.

We hypothesize that a plausible biological mechanism is the interplay between (a) the different dynamics of ON and OFF LGN cells and (b) the specific wiring that connects ON and OFF cells to V1. (b) refers here to the well-known fact that OFF and ON LGN cells are wired to segregated V1 receptive field subregions (3, 22, 23). Our main contribution is (a): we identify in **Results** dynamic differences in the ON/OFF pathways that together with (b) produce distinct response time-courses in separated receptive-field subregions. The mechanisms we propose are biologically grounded, and, as we show, they are sufficient for initiating DS in the feedforward LGN input to cortical cells.

To constrain our theory, we present novel experimental results on the responses of Macaque 4Cα Simple cells to drifting gratings. Most Simple cells we recorded in 4Cα were unambiguously DS, preferring consistently the same direction over their entire visible ranges of spatial frequency (SF) and temporal frequency (TF); about half of the cells had high DS. Our data reveal also an important characteristic of DS neurons, namely the approximate invariance of DS with SF and TF. Explaining the broad-band character of DS (in TF and SF) is a challenge for all previous theories. Our theory includes a complete analysis of how stimulus features like SF and TF interact with ON/OFF dynamics and LGN-to-cortex wiring to explain the broad-band character of DS. The theoretical predictions are in good agreement with data.

With regard to broader implications, although the theory as described in this paper is specifically about DS, an important message is that when combining information from multiple channels, slight biases in their temporal filters can greatly enhance the capability of a system. Thus, it may be possible to exploit the temporal axis further in the processing of biological and nonbiological signals, especially in the neural processing of sensory inputs and possibly in computer vision.

## Results

As discussed in the Introduction, many neurons in layer 4Cα of the Macaque are known to be direction selective (DS), but the LGN cells which provide them with feedforward inputs are not, and the challenge is to explain the origin of DS. The LGN input to layer 4Cα is sparse; the total LGN input to thousands of 4Cα cells preferring the full range of orientations and directions comes from only O(10) LGN cells (24-27). Hypothesizing that DS appears already in the summed responses of groups of LGN cells afferent to a cortical cell, we reason that mechanisms for DS likely arise from visual response properties of the LGN cells and from the way outputs of LGN cells are combined. The problem is: which LGN properties are implicated, and how do they combine to give DS? As each cortical cell receives input from two or three rows of ON and OFF cells, an idea going back to Hubel and Wiesel (3, 22), a natural candidate for the origin of DS is ON-OFF interaction. In **Results II** below we show, in the simplified setting of a pair of ON-OFF LGN cells, that DS in their summed responses can be deduced from known biological factors. These ideas are extended to more realistic LGN configurations in **Results III**. But first we present in **Results I** experimental data to document DS properties of neurons in cortical cells, setting conditions that must be met for the proposed theory of DS to follow.

### I. Experimental data

First, we show that most Simple cells in Macaque 4Cα are DS cells. The data that show it are the responses of a population of Simple cells that were recorded with microelectrodes. The visual receptive fields of the cells were located at 1-6^0^ eccentricity. For each cell, responses were recorded for high-contrast, drifting sinusoidal gratings in different orientations (ori), spatial frequencies (SF) and temporal frequencies (TF). All measurements of Simple cell responses are in terms of the amplitude of the f1 component of neural firing rate in response to the drifting grating (14, 28). We also recorded visual responses of Complex cells in layer 4Cα but they did not exhibit DS and are not considered in this paper.

In the experiments, after the optimal orientation of each cell was determined, responses to the two opposite directions at the optimal orientation were measured with what we thought were stimuli that would produce the Maximum firing rate. If response in one direction was larger, that direction was called Preferred (abbreviated as Pref) and the other was the Opposite (to Preferred, or Opp) direction. Pref and Opp directions defined by this procedure were fixed, and we studied responses in these two directions over a range of TF and SF.

Using the ratio of Pref and Opp responses as a measure of DS, we present below statistics for two groups of cells: a “highly DS” group consisting of cells with Pref/Opp > 3, and a group with Pref/Opp > 2 which we refer to as “DS”. We required Pref/Opp > 2 to ensure that the preferred direction could be meaningfully identified.

#### DS as function of TF

We have complete TF data for 53 Simple cells in layer 4Cα. Of these 53 cells, about half (25/53) were highly DS, while 75% (=40/53) were DS. The left panel of Figure 1A shows the medians of the Pref and Opp-responses for these two groups as functions of TF. Observe that for both groups, the median Pref-response is larger than median Opp-response consistently across the entire TF range. The Pref-response for the highly DS cells is temporally bandpass, with a much smaller response at 1Hz than at 10 Hz. Notice also that response amplitude drops off steeply at high TF so that at 32 Hz both Pref and Opp responses are very small and often below the noise level.

**Figure 1.**
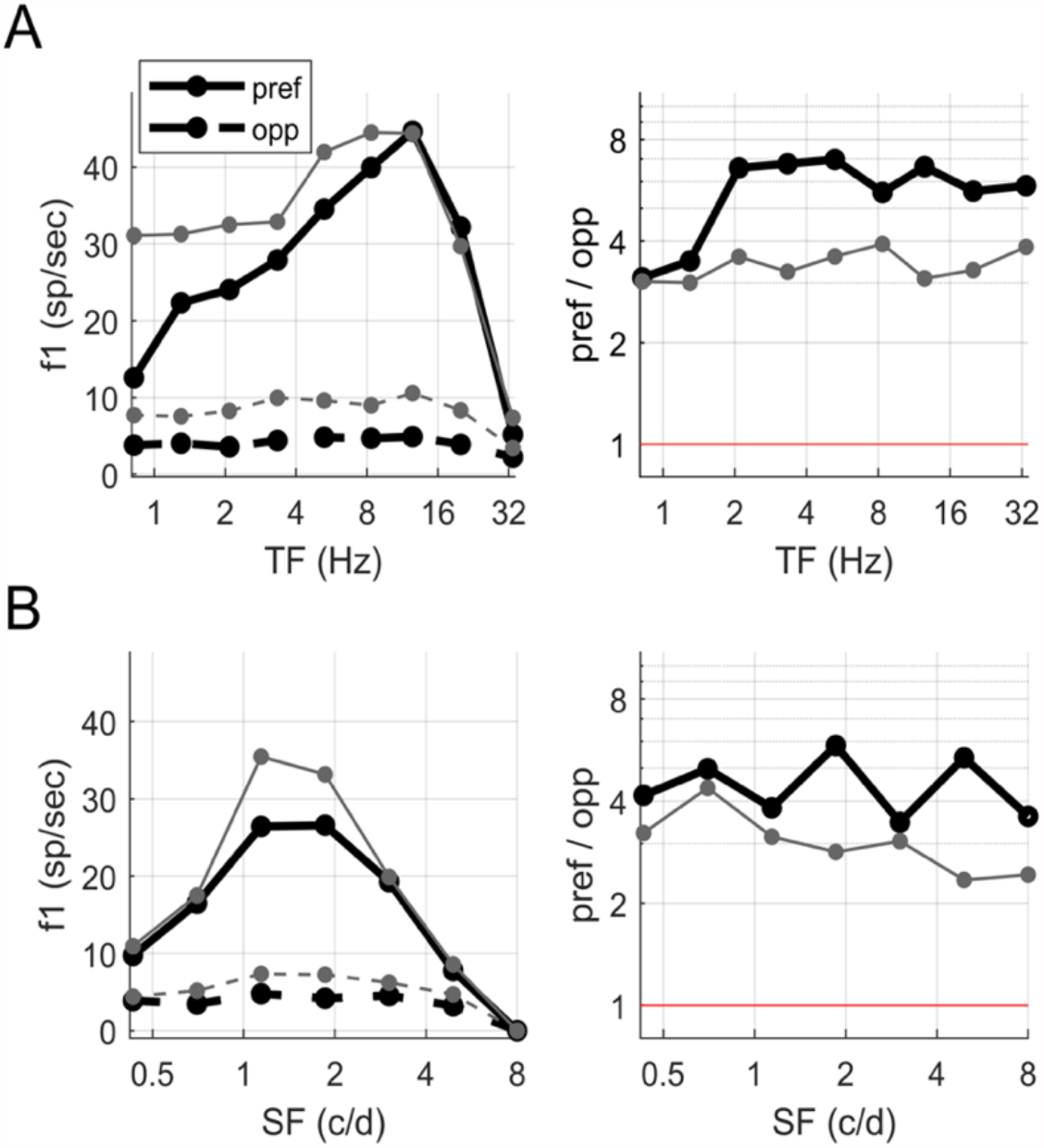
Data on firing rates and DS as functions of TF and SF. **A**. Data for 25 Simple cells with Pref/Opp > 3 (thick black) and 40 simple cells with Pref/Opp > 2 (grey) are presented. Left: Median Pref and Opp-firing rates as function of TF. Right: pref/opp as function of TF, for the same two groups of cells. Only cells for which firing rate at Max(pref,opp) > 4 spikes/s were included; below 10 Hz this condition was met by all the cells; by 32 Hz, # cells (n) dropped to 13 for Pref/Opp > 3 and to 23 for Pref/Opp > 2. Data points were far above the red line; pref/opp < 1 means directional reversal. **B**. Corresponding figures with SF in place of TF, for a collection of 26 Simple cells with Pref/Opp > 3 (thick black) and 42 simple cells with Pref/Opp > 2 (grey). Excluding low firing cells as in **A**, for Pref/Opp > 3, n = 24, 15, 3 at 3, 5, 8 c/d respectively; for Pref/Opp > 2, n = 39, 27, 8 at 3, 5, 8 c/d.

The right panel of Fig. 1A shows, for the same two groups of cells, the medians of the pref/opp ratio as functions of TF. For both groups, these ratios are roughly constant throughout the visible range of TF, demonstrating that in Macaque layer 4Cα, DS is broad-band in TF.

#### DS as function of SF

We have complete SF data for 54 Simple cells, of which 26 were highly DS and 42 were DS. Figure 1B shows the same plots as in Figure 1A but as functions of SF, where the TF was at or near optimal for the neuron. The pref/opp ratios were uniformly away from 1 throughout the visible range of SF, implying that DS in layer 4Cα neurons is broad-band in SF, though pref/opp declined gradually as SF increased from 1 to 5 c/d in the DS group.

Aligning cells on their preferred SF and TF gives similar results; see **SI**.

To summarize: Our data show that most of the Simple cells recorded in layer 4Cα had some DS, and roughly half had high DS, with Pref/Opp > 3. DS was found to be broad-band in both TF and SF; each cell had a consistent preferred direction and roughly constant pref/opp-ratios over the entire visible ranges of both TF and SF.

### II. Proposed mechanisms for DS for ON-OFF LGN pairs

**Results II** considers the summed responses of two LGN cells, one ON and one OFF. We propose mechanisms for how DS could be produced, giving a complete analysis in this simplified setting under idealized conditions. Predictions from the theory developed here are then verified in **Results III** against typical configurations of LGN afferents to cortical cells.

#### ON-OFF pair setup

We study here the sum of the responses of two LGN cells, one ON and one OFF, representing two rows of ON and OFF LGN cells. Throughout the analysis, the OFF-cell is placed on the left, the ON-cell is on the right, and they are separated spatially by d degrees, as shown in Figure 2A. The stimuli L(x,t) considered are moving sinusoidal grating patterns; most experimental data about DS in cortical neurons have been obtained with such stimuli, including those in **Results I**. Our theory generalizes to drifting bars and edges, but that will not be discussed here.

**Figure 2.**
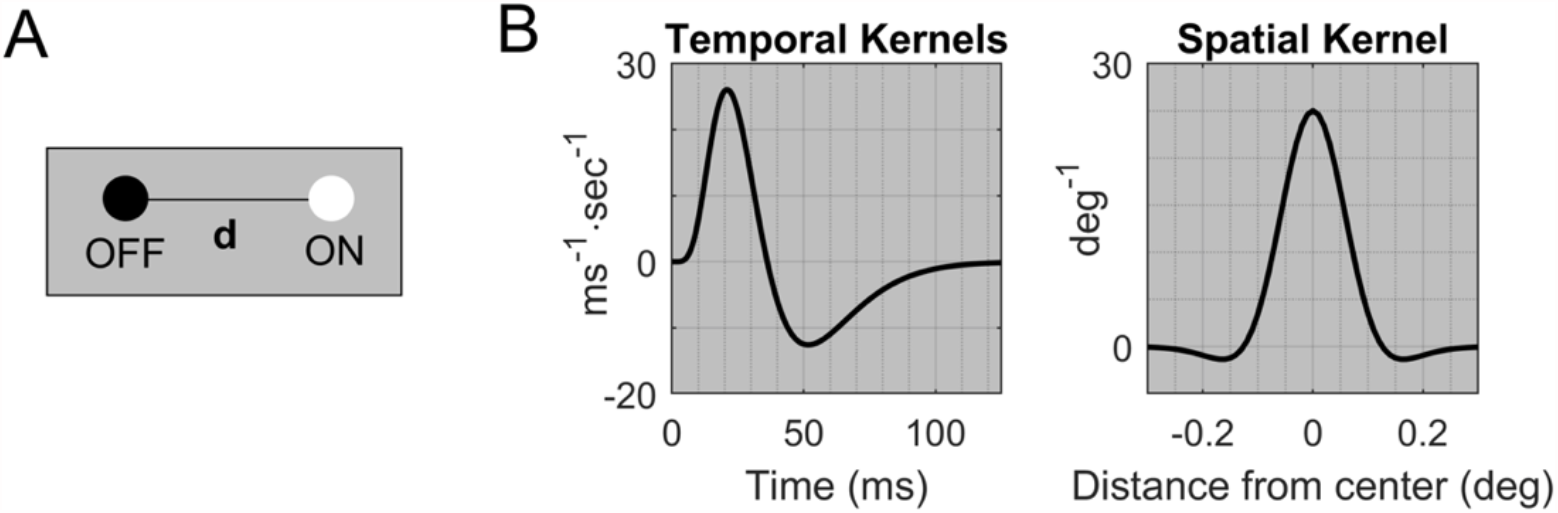
Setup for theory. **A**. A pair of ON and OFF LGN cells is separated by d°, with ON to the right of OFF. **B**. Reference temporal kernels K(t), and spatial kernel A(x,y). LGN response is assumed to be the convolution of the stimulus function with these two kernels.

We consider only gratings aligned with the rows of ON and OFF cells, so that in the simplified setting of an ON-OFF pair they are either left or right-moving. The following notation will be used: Drifting gratings are **Right** (from left to right) when L(x,t) = C sin(2π (-gx+ft)), and **Left** (from right to left) when L(x,t)=C sin(2π (gx+ft)); here g (units: c/deg) is spatial frequency (SF), f (units: Hz) is temporal frequency (TF), and C is contrast. Throughout the discussion, we assume the responses R_ON_(t) and R_OFF_(t) of the ON and OFF-cells are given by convolutions of L(x,t) against the cell’s spatial and temporal kernels, i.e.,

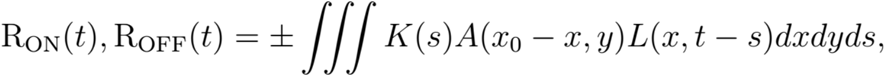

where K(s) is the temporal kernel and A(x,y) is the spatial kernel. The cells are located at (x_0_,0); x_0_ = 0 for OFF and x_0_ = d for ON. The forms of the temporal and spatial kernels are illustrated in Figure 2B; exact formulas and other details are given in **Materials and Methods**.

Of interest is R(t) = R_ON_(t) + R_OFF_(t), the sum of the ON and OFF responses.

#### DS via differential in phase differences

We start with the following questions: Does the summed response R(t) of the ON-OFF pair have a preferred direction, and what is the preferred direction? The answers are clear in the frequency domain, an insight that goes back to Watson and Ahumada (20, 21). DS depends on the phase difference between R_ON_(t) and R_OFF_(t), denoted Δφ, which takes values between 0 and π. The closer Δφ is to 0, the larger the amplitude of R(t) because then ON and OFF responses are in phase and reinforce each other. The closer Δφ is to π, the smaller is R(t)’s amplitude because then ON and OFF are in antiphase and cancel each other.

We define Δφ(→) to be the ON-OFF phase difference for a right-moving grating and Δφ(←) for motion to the left. If ON and OFF cells have the same response time-course, then Δφ(→) = Δφ(←) due to a left-right symmetry, and there is no DS (18, 20, 21). (More details are provided below, under Mechanism #1.) But if the symmetry of ON and OFF is broken and Δφ(→) ≠Δφ(←), the phase difference for one of the directions is closer to 0, and that is the preferred direction.

##### Mechanism #1. Delay of ON-response relative to OFF

A delay in the ON-response with respect to OFF is consistent with retinal circuitry required for the sign inversion in the ON pathway (29) and with direct measurements (30, 31). We consider a delay of *t*_*0*_ ms in the response of the ON-cell, and explain how that leads to unequal phase differences for the right and left-moving gratings.

To illustrate how the introduction of a delay breaks the symmetry between Δφ(→) and Δφ(←), we show in Figure 3 numerical simulations of the response of an ON-OFF pair to drifting gratings. For those simulations, f=10 Hz, around the peak TF for Magnocellular cells (12) and g=2.5 c/deg, at or near an optimal SF for many cells in V1 representing the parafovea (32) which is the region we are modeling. The distance d (Fig. 2A) between ON and OFF is set at 0.1°, a first approximation based on retinal topography (25).

**Figure 3.**
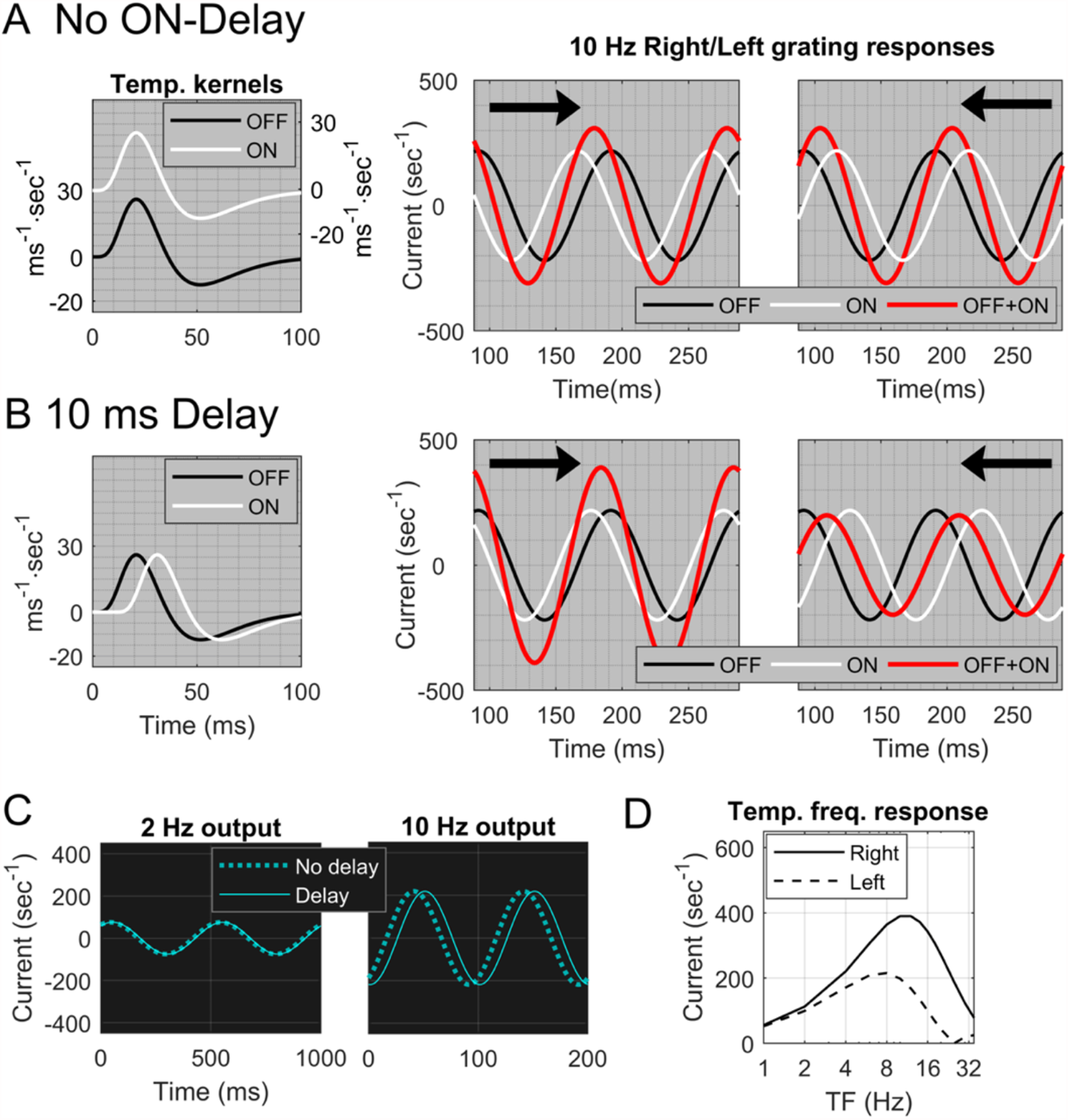
Effect on DS from time delay between ON and OFF LGN cells. **A**. Two identical temporal kernels, one OFF (black) and one ON (white) (leftmost panel). The ON kernel is displaced vertically to avoid overlap with OFF. The two panels on the right show responses of an ON-OFF pair separated by d=0.1^0^ to gratings at 10Hz and 2.5 c/deg. Waveforms of ON (white), OFF (black) and summed response ON+OFF (red), with arrows indicating the direction of motion. **B**. Two identical kernels, one OFF (black) and one ON (white), with a 10 ms delay for the ON-kernel, are shown in the leftmost panel. The two panels on the right show results analogous to those in **A**, with an ON-delay of 10 ms. **C**. The two panels show the responses of the ON-kernel with a 10 ms delay (solid) compared to no-delay (dashed) at TF=2Hz and 10 Hz respectively. **D**. Dependence on temporal frequency (TF) of the effect of delay on DS. Plotted is the F1 component of the ON+OFF sum as a function of TF for both Right (solid) and Left (dashed) directions.

In Figure 3A, there is zero delay. ON and OFF responses are shown separately (in white and black), and their sum is shown in red. The responses to left and right motion are identical with the roles of ON and OFF interchanged, resulting in the same summed response amplitude for Left and Right. In Figure 3B, a delay *t*_*0*_ between ON and OFF is introduced (left panel). Data (30, 31) suggest that *t*_*0*_ is likely between 5-15 ms. Here we use *t*_*0*_ = 10 ms. The summed response R(t) is larger for Right motion, for which the ON and OFF responses are closer to being in-phase (Figure 3B); Right is the preferred direction.

To understand the effect of the delay, we analyze ON/OFF phase differences in the Right and Left directions. For Right, ON lags OFF because of the separation d and the delay t_0_; there is also a phase difference of π (=180°) because of the sign difference between ON and OFF. Thus, Δφ(→) = [2 π (-gd -ft_0_) + π] where we use the formalism that [x] denotes the shortest distance between x and the set {0, ±2 π, ±4 π, …}. (The phase difference is distance between two points on a circle and all multiples of 2 π are equivalent to 0.) For Left, Δφ(←)= [2 π (gd -ft_0_) + π]; in the Left direction, the ON cell has a phase lead due to the separation d, but a phase lag due to the delay. Response amplitudes in the two directions are predictable from the phase differences. In Figure 3B, Δφ(→) is smaller so Right is the preferred direction. The formulas above also show that when t_0_=0, Δφ(→) = [-2 π gd+ π] =[2 π gd+ π] = Δφ(→), hence there is no preferred direction.

Because the delay enters Δφ as *2 π ft*_*0*_, ON time-delays are effective in producing DS at higher TF, but not at lower TF (Figure 3D). For instance, a grating with TF=10 Hz travels 1/10 of a cycle in 10 ms, whereas at TF=2 Hz, 10 ms is only 1/50 of a cycle. Figure 3C shows the phase shifts of an ON-cell due to a 10-ms delay in response to gratings with TF = 2 and 10Hz. The 10Hz grating produced significant DS (Figure 3B). However, for the 2Hz grating, the phase shift is negligible and the picture is closer to that of no-delay; it does not produce much DS (Figure 3D).

The data in Figure 1A show, however, that DS is present both at high and low TF in the Macaque. We therefore need another mechanism in addition to the time-delay.

##### Mechanism #2. Modified structure of ON-kernels

Another suggestion from LGN data (30) is that the waveforms of ON and OFF time-kernels have different temporal structure. We use the following notation: Let the OFF-kernel *K*_*OFF*_*(t)* = K(t) where K(t) is as in Figure 2B (see **Materials and Methods** for more detail), and say a temporal kernel *K*_*a,b*_*(t)* has structure (a,b) if

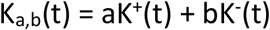

where K^+^(t) = max{0, K(t)} and K^-^(t) = min{K(t), 0} are the positive and negative parts of the function K(t) respectively. In this notation, *K*_*OFF*_*(t)* has the structure (1,1). It can be shown mathematically (see **Materials and Methods**) that a kernel *K*_*a,b*_*(t)* with *a≠b* produces a phase shift relative to *K*_*1,1*_*(t)*. For *a>b*, it is a <u>phase lag</u>, and the lag is more prominent at low TF and declines as TF increases (Figure 4A). Other kernels *K*_*a,b*_*(t)* with *a>b* produce similar results. In general, the larger a/b, the larger the shift. Ratios of a/b ∼ 2 are consistent with data for ON-kernels (30).

**Figure 4.**
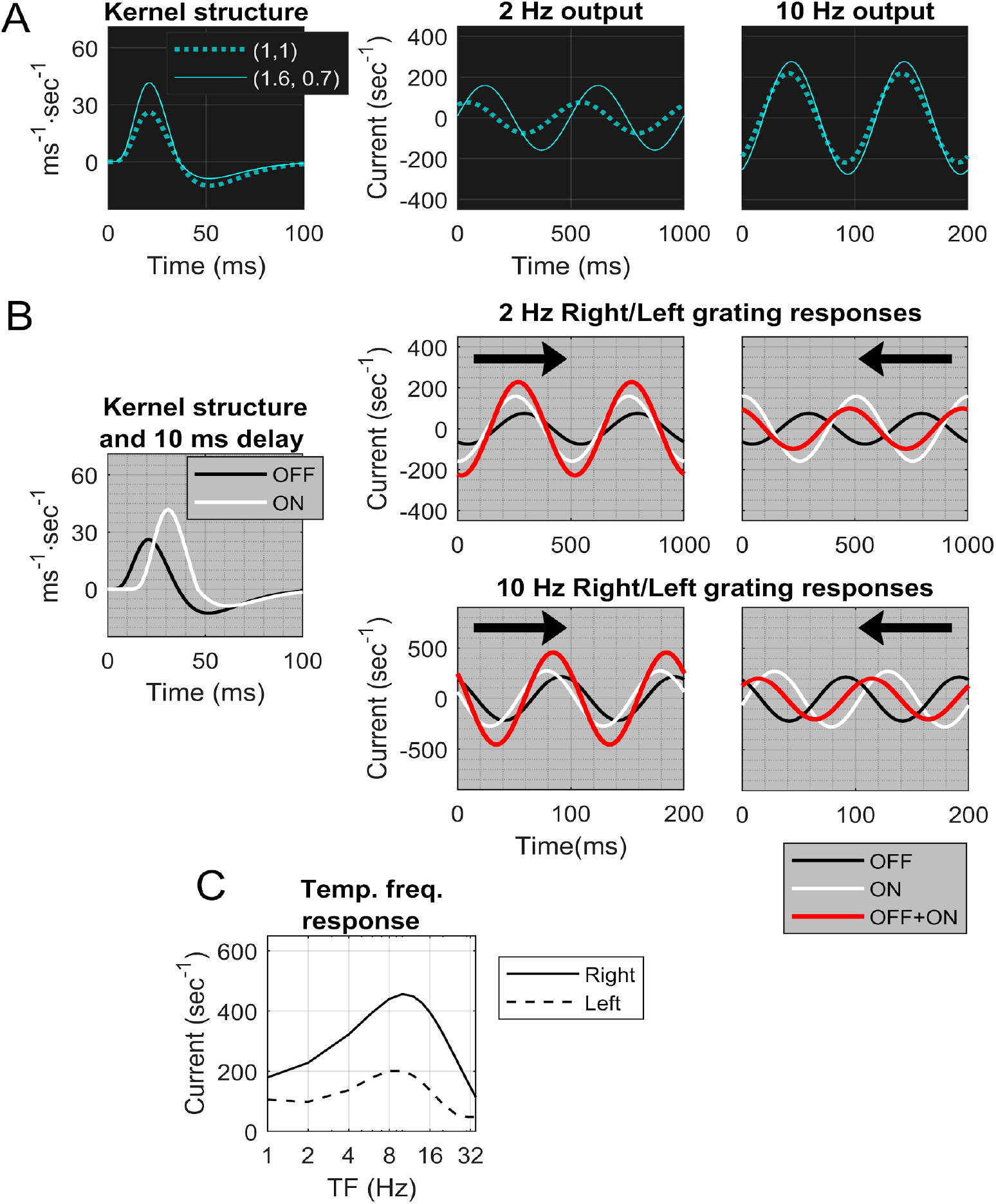
Effect on DS of time delay and modified temporal kernel structure. **A**. The left panel shows a kernel of type (1.6, 0.7) (solid) and one of type (1,1) (dashed). The two panels on the right show the phase lags of the (1.6, 0.7)-kernel (solid) relative to one of type (1,1) (dashed) at TF = 2 Hz and 10 Hz. **B**. The setup is like that of Figure 1; d = 0.1^0^ and SF=2.5 c/deg. The ON-kernel is of type (1.6,0.7) in addition to having a 10 ms delay with respect to OFF. In the righthand panels the ON responses (white) and OFF responses (black) have greater phase differences for Left than for Right. Summed waveforms (red) in Right(-->) and Left(<--) directions for TF=2Hz and TF=10 Hz reveal Right preference at both TFs. **C**. Right vs Left responses as functions of TF, at SF=2.5 c/d. It is the analogue of Fig 1D but with a (1.6, 0.7) ON-kernel in addition to the 10 ms delay.

Combining Mechanisms #1 and 2, i.e., letting *K*_*OFF*_*(t)= K*_*1,1*_*(t)*, and letting the ON-kernel *K*_*ON*_*(t)= K*_*a,b*_*(t)* for some a>b *with* a delay of *t*_*0*_ ms, we may write Δφ (→)and Δφ(←) as

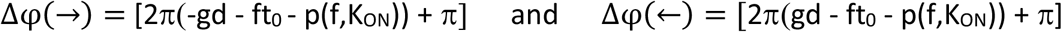

where p(f,K_ON_)>0 is the phase lag contributed by the temporal structure of the ON kernel, and [] has the same meaning as before.

From the discussion above, it follows that Mechanisms #1 and 2 combined would produce DS in the summed response of an ON-OFF pair across a wide range of TF with a consistent preferred direction. Simulations confirm the correctness of this reasoning. Figure 4B shows the individual and summed responses for left and right-moving gratings, where the ON-kernel has temporal structure (1.6, 0.7) in addition to a time delay of 10 ms, for both f=2Hz and f=10Hz. The response shows a clear preference for Right motion.

Figure 4C confirms these results across the full range of TF. We compare this panel to Figure 1A: Figure 4C is our theoretical prediction of the summed *feedforward inputs* to a cortical cell (for left and right gratings), while Figure 1A, left, shows data on the *responses* of DS cortical cells. Similarities in the shapes of the input and output curves as functions of TF and the fact that pref/opp for both are roughly constant over the entire range of TF (Figures 4C and 1A, right) lend credibility to the theory proposed.

#### Preferred direction depends on d and SF (equivalently RF size)

In Figures 3 and 4, the preferred direction is from OFF to ON (Right), but is that always the case? If not, then is there a direction that is consistently preferred by an ON-OFF pair, or does the preference vary with stimulus? To answer these questions systematically involves examining the relation between the spatial frequency (SF) of the grating and the physical separation d between the ON and OFF pair.

A crucial observation is that if we fix d and vary SF, then the preferred direction switches at gd = 1/2, 1, 3/2 … where d is ON/OFF separation and g is SF. Specifically, in our OFF-ON configuration with OFF on the left, when gd < 1/2, i.e., when d < 1/2 the spatial cycle of the grating, Right motion is preferred (this was the case in the examples shown in Figures 3 and 4). For 1/2 < gd < 1, Left motion is preferred, and directional preference is reversed again as gd increases past 1. We found that preferred directions are determined solely by the relation between d and SF, and are independent of the TF of the grating, provided that the temporal shift ft_0_ + p(f,K_ON_) is < π, a condition easily satisfied by the mechanisms proposed. A mathematical proof for the assertions in this paragraph is given in **Materials and Methods**.

Figure 5A illustrates the consequences of these predictions by showing Right and Left responses as functions of SF, for four different values of d, and five TFs. In the rows in which d=0.15 or d = 0.2 deg, the predicted directional switch can be seen as SF increases in each plot: When g < 1/(2d), the preferred direction is from OFF to ON (Right), and when g > 1/(2d), it is from ON to OFF (Left), and this is true across the entire range of TF. Reversals for d=0.1 are less visible because ON/OFF separation of d=0.1° corresponds to reversals at 5 c/d, where the curves have steep slopes. As for d=0.05°, reversals are predicted to occur at 10 c/d, a high enough SF that the response is too small to be visible.

**Figure 5.**
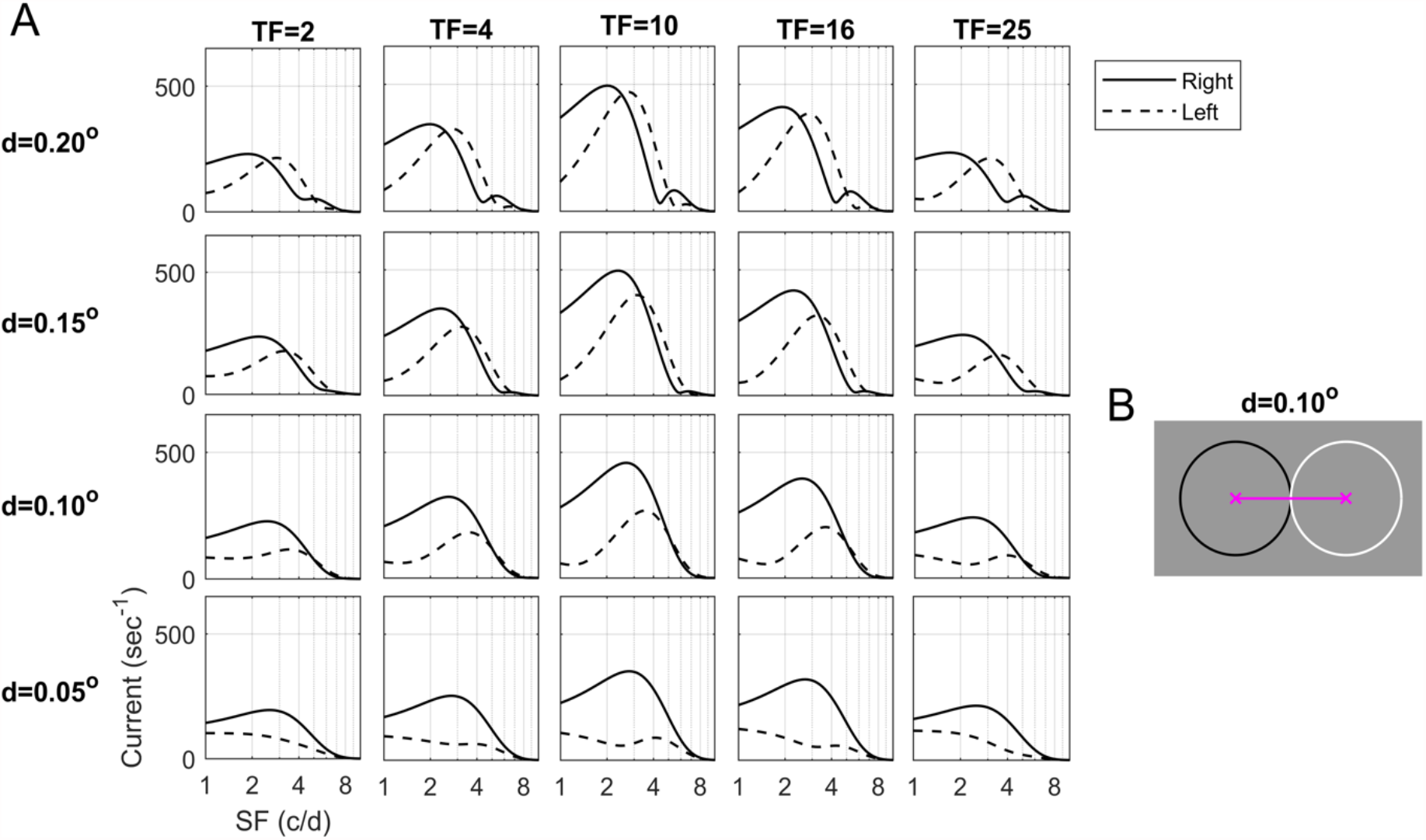
Dependence of the DS of the ON-OFF pair on SF and d. **A**. Matrix of response vs SF plots is drawn parametric in d and TF. Rightward and Leftward responses are plotted vs TF for 4 different values of d: 0.05, 0.1, 0.15, 0.20^0^. **B**. Two LGN cells separated by d=0.1°. The circles are 1 SD of their spatial kernels.

The following points are illustrated in Figure 5A. One is that while, mathematically, direction reversals occur at gd = 1/2, 1, 3/2 …, as a visual response a reversal is registered only if the SF at which it occurs falls within the animal’s range of visible SF, which is known to scale with rf size (32). A second point is that as important as the phenomenon of reversal itself is the loss of DS near reversal points. The lowering of Pref/Opp for an entire region of (d, SF) near reversal occurs as a result of the gradual interchange of preferred directions.

That leads to our third point, the prediction that DS is stronger for smaller ON/OFF separation. In Figure 5A, peak firing rate occurs at TF=10 Hz and SF = 2-3 c/d, a good approximation to data (Figure 1). Observe that at these values of TF and SF, Pref/Opp is close to 1 at d=0.2°, a little larger at d=0.15°, and continues to increase as d gets smaller. The Pref/Opp ratio is larger for smaller d because, with decreasing d, reversal occurs farther from SF=2-3 c/d.

#### Implications for the Macaque

In Macaque V1, we have not observed evidence for reversals of Preferred Direction (Figure 1B, right), suggesting that reversal occurs outside of the range of SF that produces a response. This range can be seen in Figure 1B, left. The discussion above together with the results in Figure 1 predict therefore that to provide feedforward input with robust DS across the full range of SF, ON-OFF rows that project to the same cortical cell are likely < 0.1° apart at the eccentricities of the cells recorded. The decline in DS for SF between 1 and 5 c/d seen in Figure 1B suggests also that we may be heading for a reversal point, so this ON-OFF separation is likely > 0.05°.

Comparing theory to data, our theoretical prediction of the summed LGN feedforward inputs to cortical cells at TF=10 Hz (the optimal TF) and d = 0.05-0.1° as shown in Figure 5 shows a remarkable resemblance to the data on cortical responses shown in Figure 1B, left.

### III. DS in feedforward LGN inputs

In **Results II**, we developed a theory for the DS of an ON-OFF pair. We now extend this theory to typical feedforward LGN inputs to layer 4Cα, assuming that the set of LGN cells that project to a simple cell in layer 4Cα is organized in two or three rows of ON and OFF cells, alternating in ON/OFF in the case of three rows. These widely accepted rules of LGN-to-cortex connections were proposed by Hubel and Wiesel and have been confirmed in experiments (23, 34-36). The setup is as described at the beginning of **Results II**, but for arbitrary collections of LGN cells. The response of each LGN cell is computed as a convolution of the light intensity function with spatial and temporal kernels as before, and DS for the summed responses from the entire group is evaluated.

LGN sampling of visual space is known to be very sparse; two independent estimates based on M-retinal ganglion cell density (25) and on Macaque LGN data (24) point to an average of about 9 LGN cells projecting to an area of 0.25° x 0.25° in the visual field at 5° eccentricity. Respecting this density, we modeled a sheet of LGN cells using either a randomly perturbed hexagonal lattice (Figure 6 A-C) or a randomly perturbed rectangular lattice (Figure 6 D-E). Once the ON and OFF mosaics were fixed, the temporal kernel of each LGN cell was drawn randomly and independently following the prescription from **Results II**.

**Figure 6.**
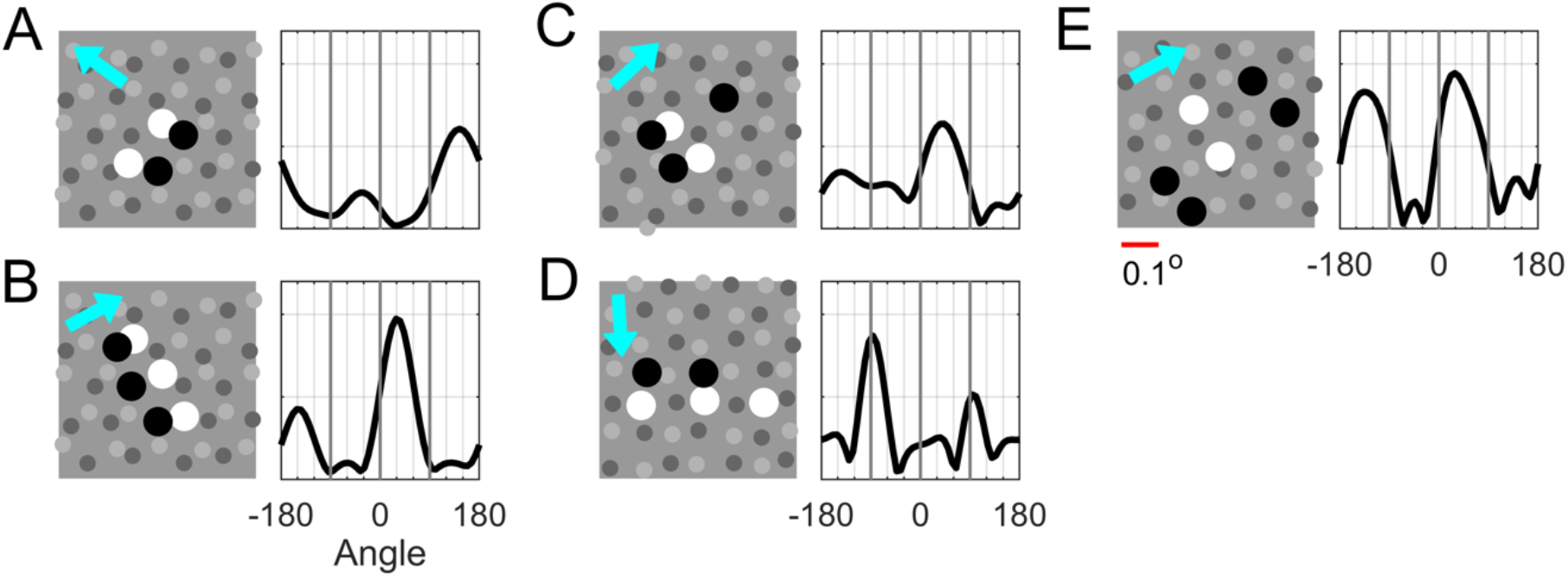
Sample LGN configurations and orientation (ori) tuning of their summed responses. Shown are 5 examples (**A-E**) of configurations of LGN cells that project to a simple cell in layer 4Cα (left) and ori tuning curves of their summed responses (right). Grey dots indicate locations of LGN cells present: ON in lighter grey, OFF in darker grey. Larger black and white dots are the OFF and ON cells in the configuration. LGN kernels are randomly drawn, and are of type (a,b): for OFF, a,b ∈ (0.9,1.1), and for ON, a ∈ (1,2) with a/b ∈ (1.4, 2.2); ON-delay time ∈ (9,11) ms, also randomly drawn. In ori tuning curves for summed responses, cyan arrow indicates preferred direction according to tuning curves. Scale bar of deg shown below Panel E.

We consider in Figure 6 some typical configurations of LGN cells that form the inputs to a cortical cell. Following (37), it is likely that no more than 4-6 LGN cells project to a Simple cell, and we learned from **Results II** that ON/OFF separation d should be < 0.1o, with stronger DS predicted for smaller.

The configurations in Panels A and B translate easily to the ON-OFF setting studied in **Results II:** projecting along a direction orthogonal to the rows of ON and OFF cells, we arrive at precisely the picture treated in **Results II** with near-optimal spacing between the two cells. As expected, DS of the summed responses of these configurations are strong, and as predicted, the preferred direction is from OFF to ON.

Panels C and D require extensions of the ideas developed for ON-OFF pairs in **Results II**. In Panel C, the projected configuration has three cells, an ON-cell flanked by two OFFs. DS for this 3-cell configuration can be thought of as competition for DS-dominance between the OFF-ON pair on the left and the ON-OFF pair on the right; one should also expect cancellation as these two pairs will prefer opposite directions. In the configuration shown in Panel C, one would predict that the left OFF-ON pair should prevail, because it carries more weight and ON/OFF distance is smaller. The orientation tuning curve shown confirms these predictions: the left ON-OFF pair is dominant, and DS is weaker due to partial cancellation. Panel D projects to an ON-OFF pair, with the ON-cell carrying significantly more weight. That weakens the DS, though the preferred direction is still quite evident.

The configuration in Panel E projects to an OFF-ON-OFF triplet that is left-right symmetric; no DS can be expected for such a configuration and little is observed.

Figure 6 and other examples not shown illustrate the fact that the mechanisms proposed in **Results II** lead to variable amounts of DS in feedforward LGN inputs depending on the spatial alignment of the group of LGN cells (and their temporal kernels). The conclusion about variable amounts of DS is consistent with the data presented in **Results I**, which show that not all 4Cα neurons are DS, and among those that are DS, Pref/Opp can take on a range of values; see Figure 1.

## Discussion

The main contributions of this paper can be summarized as follows:

1. We presented new experimental data that document (i) the abundance of high DS cells in layer 4Cα of Macaque V1, and (ii) the dependence of DS on TF and SF for these cells. We then proposed a theory the predictions of which are consistent with these data.
2. Our theory asserts that DS could originate from dynamic differences between ON and OFF cells and should therefore already present in feedforward LGN inputs to cortex. Two dynamic differences, time delay and kernel shape, are sufficient for initiating DS across the full spectrum of TF.
3. We made precise how stimulus features such as TF and SF, the dynamics of ON/OFF LGN cells, and LGN-to-cortex wiring interact to determine the preferred direction and magnitude of DS in feedforward LGN inputs.

These results should be understood in the following light:

First, **Results II** discusses mechanisms for DS under idealized conditions. Not all groups of LGN cells afferent to a cortical cell can be projected neatly to an ON-OFF pair, as we have explained in **Results III**. Time delay for ON-cells in Mechanism #1 can, in reality, take on a range of values, and the kernels for ON-cells in Mechanism #2 are not always of type (a,b) with a ∼ 2b as suggested by the examples in (30). We expect data in relation to the properties in Mechanisms #1 and 2 to be more variable than in the theory described in **Results II**, consistent with the fact that not all cells in 4Cα have high DS (see Figure 1).

Second, we do not claim that the mechanisms proposed here are the only mechanisms that contribute to DS. The theory is about DS properties in feedforward LGN inputs to cortex, and it asserts only that DS could *originate* from dynamic differences between ON and OFF LGN cells. Cortico-cortical interaction can modify DS properties. We will report on that in a separate paper in which we implement the ideas of the present paper to build a large-scale cortical model (38). It will be shown in (38) that Mechanisms #1 and 2 alone are sufficient for producing a degree of DS comparable to that found in data. The presence of specific intra-cortical connections would not contradict our theory, however, especially in layers beyond 4Cα where longer-range connections are present; see e.g. (13).

Previous theoretical work has proposed spatio-temporal inseparability (STI) in x-t plots as a characterization of DS (18,20, 21). Findings of STI have been obtained for V1 neurons in experimental studies (39, 40). While x-t plots in and of themselves are not intended to shed light on mechanisms, STI is an indication of DS. In **Supplemental Information (SI)** we show that when the mechanisms proposed in **Results II** are implemented, the summed responses of ON-OFF pairs of LGN cells generate STI. The converse need not be true, however: STI need not imply DS. The amount of DS depends also on the spatial and temporal properties of the visual stimulus, a fact we also demonstrate in **SI**.

We finish with comparisons of our theory for DS in Macaque layer 4Cα with theories of DS offered for other species. One neural mechanism for DS proposed in previous modeling studies was for specific connections between V1 cells and LGN cells that are either Sustained or Transient in their step responses (41, 42). The Sustained-Transient theory was proposed originally as a mechanism for DS in cat V1 cortex. The concept was revived recently in models of DS in mouse V1, by Lien and Scanziani (14) and Billeh et al (15). We consider the Sustained-Transient theory to be implausible for Macaque because there are not clearly separate classes of Sustained and Transient Magnocellular LGN neurons. In our proposed theory, ON vs OFF LGN cells have different temporal kernels, but these differences are much smaller and subtler than those observed in Sustained and Transient neurons in rodent or cat retinal and LGN cells (16). Even accepting Sustained/Transient differences as underlying mechanisms of DS for cat and mouse V1, the story there is incomplete without information on the wiring of these two subpopulations and how that leads to the DS properties observed. In the mouse, estimates are that there are either ten (14) or forty (15) LGN cells converging into one subfield. The problem of how to place all (or most) of the Sustained inputs into one subfield and Transient inputs into the other remains to be clarified. Our theory solves this problem by postulating that ON and OFF cells have different temporal impulse responses. That together with the consensus view on how to segregate ON and OFF inputs from LGN into different subfields (22, 23) is sufficient for generating DS.

A different mechanism for DS is the idea that there are different types of LGN cells, so-called lagged and non-lagged cells, with widely different temporal kernels. This mechanism has been invoked to explain DS data in cat (19) and ferret (43) visual cortical cells, and to be the source of STI in the motion-energy model. It is important to note that the data on the TF dependence of DS in ferret V1 (43) are very different from the Macaque data in Figure 1, suggesting that the temporal dependence of cortical DS in ferret and cat may be very different from that in Macaque. The lagged vs. non-lagged theory does not predict the broad-band character of DS found in Macaque 4Cα cells; it predicts that DS should weaken markedly at high TF (43) as in the ferret V1 data. Therefore, the mechanism of DS in Macaque V1 must be sought elsewhere. In support of that conclusion, lagged cells have not been found in Macaque LGN (9-12).

Other DS models invoked intra-cortical inhibitory mechanisms for DS in cat V1 (16, 44, 45), in particular cortical inhibition tuned to the non-preferred direction, but intracellular-recording in cat V1 (40) showed such models also to be implausible. Our theory indicates that direction-tuned cortical inhibition is not necessary for the initiation of DS in primate V1.

## Materials and Methods

### Modeling LGN response

We model the input current to an LGN cell the center of whose receptive field is located at (*x*_*0*_, *y*_*0*_) by

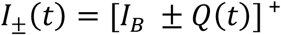

where

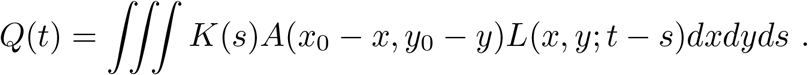

Here the ± sign is for ON and OFF-cells respectively, *I*_*$*_ is background current, *K(s)* and *A(x,y)* are the temporal and spatial kernels of the LGN cell, *L(x,y;t)* is the light intensity map, and *C* is a constant to be adjusted, and []^+^ denotes the maximum of the bracketed value and 0.

In **Results II**, the OFF cell is placed at *(x*_*0*_, *y*_*0*_*) = (0,0)*, the ON cell is placed at *(x*_*0*_, *y*_*0*_*) = (d, 0)*,and L(x,y;t) depends only on x and t. The more general formulas above are used in **Results III**. In both **Results II and III**, we have used *±Q(t)* as our ON/OFF LGN *response*, conflating the input current to LGN with its output (in addition to omitting the background firing and the []^+^ part of the operation). This conflation was on purpose: the two quantities are roughly proportional, and the exact sinusoidal form of *±Q(t)* makes the analysis much more transparent.

Formulas for the spatial and temporal kernels are as follows: The spatial kernel is a difference of Gaussians

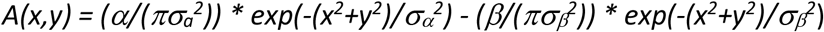

With *α*=1.0, *β*=0.74, *σ*_*α*_=0.0894, *σ*_*β*_ =0.1259 (42); *σ*_*α*_ and *σ*_*β*_ are chosen to make the center part of the receptive field correspond to a Gaussian with std. dev. ∼ 0.05 deg (13). The preferred spatial frequency of the spatial kernel *A(x,y)* is 2.5 c/deg. The reference temporal kernel K(t), also referred to as K1,1(t) in the main text, has the form (adapted from Zhu et al (46))

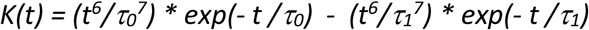

with τ_0_ = 3.66 ms and τ_1_ = 7.16 ms. Parameters are chosen following guidance from temporal kernels derived in experiments (30); specifically the cross-over point from positive to negative was chosen to lie at 36 ms so that the peak response occurs at about TF=10 Hz following the data reported in Figure 1A, left. Note that K(t) is positive before the zero crossing, as the polarity of the LGN cell is implemented in the sign of +/-in the expression for I(t), not in the kernel.

### Theoretical basis for Mechanism #2

Mechanism #2 is based on the observation that at low TF, the response to a sinusoidal signal is effectively a cosine function for (1,1)-kernels, and modifying it to an (a,1)-kernel with a≠1 adds a sine function to the response, causing a phase displacement.

In more detail, we consider a signal of the form *sin(2πft)*. For a>1, let R_1_(t) and R_a_(t) denote the responses of an LGN cell with a (1,1)-and (a,1)-kernel respectively. If the (1,1)-kernel is K(t), then the (a,1)-kernel is K(t) + (a-1) max{K(t), 0}. At low TF, convolving a function with a (1,1)-kernel is similar to computing its derivative – without dividing by the distance between the centers of mass of the positive and negative parts of the (1,1)-kernel (here that distance is ∼ 0.04 s). Convolving with a nonnegative kernel is simply averaging. Assuming ∫ max{K, 0} = – ∫ min{K,0} =1, we obtain

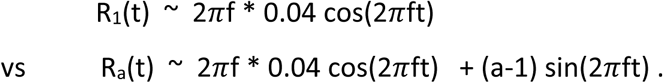

For example, for a 2 Hz grating, 2πf * 0.04 ∼ 0.5. For a (2,1)-kernel, this gives a phase difference ∼ π/3 between R_1_ and R_2_. The smaller TF, the larger the phase displacement caused by the sine term, providing DS at low TF when a simple ON-delay (Mechanism #1) does not. (For larger TF, this argument loses its validity when kernel widths are as large as temporal cycles.)

It remains to explain why an (a,1)-kernel with a>1 causes a delayed response in the ON-cell. This is because the maximum of R_1_(t) occurs at t = 0. For a>1, R’_a_(0) > 0, i.e., R_a_(t) is still increasing when R_1_(t) is at its maximum, hence R_a_(t) peaks later, as illustrated in Fig 4a.

### Mathematical analysis of reversal of directional preference

In **Results II**, we asserted that preferred direction switches at gd = 1/2, 1, 3/2 … where d is ON/OFF separation and g is the SF of the grating. A mathematical proof goes as follows. Look at the quantities inside the brackets [] in the formal expression for phase differences derived in **Results II**, and recall that without the temporal phase lag, the spatial phase lead for ON is -2πgd + π for Right-moving gratings and 2πgd + π for Left-moving gratings. Consider first the case gd < ½ (Figure 7A), where -2πgd + π is between 0 and π, and 2πgd + π is between π and 2π. These phase leads are depicted as dashed red arrows. The effect of the temporal phase lag for ON is to shift these arrows to the left, to the solid red arrows depicting the net phase leads for ON. Provided that the temporal phase shift ft_0_+p(f) is less than π (the mechanisms proposed produce shifts < π/2), we have Δφ (→) < Δφ (←), because the red arrow for Right is closer to 0 than the red arrow for Left is to 2π Hence Right motion is preferred.

**Figure 7.**
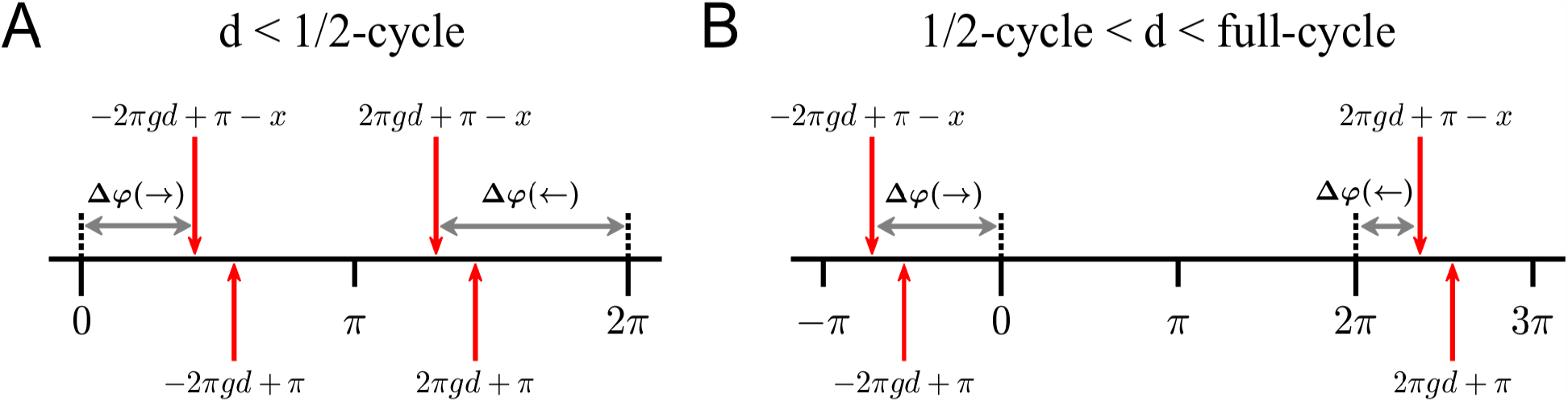
Mathematical analysis of interplay between d and SF. **A**. Phase diagram when separation d < 1/2 cycle of the stimulus grating. Dashed red arrows are spatial phase leads for ON without taking into account temporal phase lags; solid red arrows take into temporal phase lags. Red arrows between 0 and π are for Right gratings; arrows between π and 2π are for Left. The quantity *x* = ft_0_+p(f,K_ON_) is temporal phase shift, and grey lines with arrowheads are *Δφ(→)* and *Δφ(←)*. **B**. Phase diagram when 1/2-cycle < d < full cycle.

The picture is different for ½ < gd < 1 (Figure 7B). For this range of d, -2πgd + π lies between -π and 0, and 2πgd + π lies between 2π and 3π. Here the temporal phase lag causes Δφ(→) > Δφ(←), resulting in a preference for Left motion.

A similar argument shows that preference is reversed again at gd = 1.

### Experimental Methods

#### Animal Preparation

Adult male old-world monkeys (M. fascicularis) were used in acute experiments in compliance with National Institutes of Health and New York University Animal Use Committee regulations. The animal preparation and recording were performed as described in detail previously (47, 48). Anesthesia was initially induced using ketamine (5-20 mg/kg, i.m.) and was maintained with isofluorane (1-3%) during venous cannulation and intubation. For the remainder of the surgery and recording, anesthesia maintained with sufentanil citrate (6-18 µg/kg/h, i.v.). After surgery was completed, muscle paralysis was induced and maintained with vecuronium bromide (Norcuron, 0.1 mg/kg/h, i.v.) and anesthetic state was assessed by continuously monitoring the animals’ heart rate, EKG, blood pressure, expired CO_2_, and EEG.

After the completion of each electrode penetration, 3–5 small electrolytic lesions (3 μA for 3 s) were made at separate locations along the electrode track. At the end of the experiments, the animals were deeply anesthetized with sodium pentobarbital (60 mg/kg, i.v.) and transcardially exsanguinated with heparinized lactated Ringer’s solution, followed by 4 L of chilled fresh 4% paraformaldehyde in 0.1 M phosphate buffer, pH 7.4. The electrolytic lesions were located in the fixed tissue and electrode tracks were reconstructed to assign the recorded neurons to cortical layers as described previously (8). The data from the current study were obtained during the course of the basic characterization of neuronal receptive field properties that is standard for many visual cortical experiments. In 49 animals, more than 700 neurons were recorded in oblique penetrations. For this paper, a full data set was obtained for 94 neurons localized to layer 4Cα.

#### Characterization of visual properties of V1 Neurons

We recorded action potentials (spikes) extracellularly from single units in V1 using glass-coated tungsten microelectrodes. Each single neuron was stimulated monocularly through the dominant eye (the non-dominant eye occluded). Receptive fields were located in the visual field between 1 and 6^0^ from the center of gaze. Stimuli were displayed at a screen resolution of 1024×768 pixels, a refresh rate of 100 Hz, and a viewing distance of 115 cm on either a Sony Trinitron GDM-F520 CRT monitor (mean luminance 90-100 cd/m^2^) or an Iiyama HM204DT-A CRT monitor (mean luminance 60 cd/m^2^). Monitor luminance was calibrated using a Photo Research PR-650 spectroradiometer and linearized via a lookup table.

The response to drifting gratings was used to characterize visual response properties. In this paper, we report measurements of orientation tuning, spatial frequency (SF) and temporal frequency (TF) tuning. We used the f1 response for neurons (traditionally called simple cells) with f1/f0 ratios > 1.

#### Orientation tuning

The responses of each neuron were recorded to different orientations between 0 – 360^0^, either in 20 or 15^0^ steps. The stimuli were achromatic gratings at the preferred SF and TF, at a contrast of 64% or greater. All stimuli were presented in a circular window confined to the classical receptive field (CRF).

#### SF tuning

Each neuron was presented with a range of spatial frequencies, usually in ½ octave steps from 0.1 c/deg to around 10 c/deg. For some neurons, the upper limit was extended if the neurons had responses to higher spatial frequencies. For almost all neurons that were orientation selective we measured SF tuning at the preferred orientation and drift-direction as well as at the non-preferred drift direction.

#### TF tuning

Each neuron was presented with a range of temporal frequencies, usually in 1 octave steps from 0.5 Hz to 32 Hz. Measurements were made for drifting gratings at the optimal orientation and in the preferred and non-preferred directions.

## Acknowledgements

This research was supported by R01 EY001472 (RMS), NIH R01 EY008300 (MJH) and NSF 1734854 (RMS & LSY). Thanks to the graduate students and postdoctoral fellows who participated in the experiments: Dario Ringach, Michael Sceniak, Elizabeth Johnson, Siddhartha Joshi, J.A. Henrie, Patrick Williams, Dajun Xing, Christopher Henry and Anita Disney; and to Madhura Joglekar for early numerical explorations

## References

1. Parker, A.J. & Newsome, W.T. Sense and the single neuron, probing the physiology of perception. Annu. Rev. Neurosci. 21, 227–77 (1998).

2. Newsome W.T., Wurtz R.H., Dursteler M.R., & Mikami, A. Deficits in visual motion processing following ibotenic acid lesions of the middle temporal visual area of the macaque monkey. J. Neurosci. 5(3), 825–840 (1985).

3. Hubel, D.H. & Wiesel, T.N. Receptive fields, binocular interaction and functional architecture in the cat’s visual cortex. J. Physiol. 160(1), 106–154 (1962)

4. Zeki, S.M. Functional specialization in the visual cortex of the rhesus monkey. Nature 274, 423–428 (1978).

5. Movshon, J. A., Adelson, E.H., Gizzi, M.S., & Newsome, W. T. The analysis of moving visual patterns, in Study Week on Pattern Recognition Mechanisms: April 25 - 29, 1983, Volume 54 of Pontificiae Academiae Scientiarum scripta varia (ed. Chagas C., Gattass R., Gross C.) 117–151 (Pontificia Acad. Scientiarum, Rome, 1985).

6. Kiper, D.C., Levitt, J.B., Gegenfurtner, K.R. Extrastriate visual areas, in Color Vision, from genes to perception Chapter 13 (ed. Gegenfurtner, K.R. & Sharpe, L.B.) (Cambridge Univ Press, Cambridge UK, 2001).

7. Pack, C.C., Born, R.T., Livingstone, M.S. Two-dimensional substructure of stereo and motion interactions in macaque visual cortex. Neuron 37, 525–35 (2003).

8. Hawken, M.J., Parker, A.J., & Lund, J.S. Laminar organization and contrast sensitivity of direction-selective cells in the striate cortex of the old-world monkey. J. Neurosci. 8, 3541–3548 (1988).

9. Wiesel, T.N. & Hubel, D.H. Spatial and chromatic interactions in the lateral geniculate body of the rhesus monkey. J. Neurophysiol. 29, 1115–1156 (1966).

10. Kaplan, E. & Shapley, R. X and Y cells in the lateral geniculate nucleus of the macaque monkey. J. Physiol. 330, 125–143 (1982).

11. Hicks, T.P., Lee, B.B., & Vidyasagar, T.R. The responses of cells in macaque lateral geniculate nucleus to sinusoidal gratings. J. Physiol. 337, 183–200 (1983).

12. Derrington, A.M. & Lennie, P. Spatial and temporal contrast sensitivities of neurons in lateral geniculate nucleus of macaque. J. Physiol. 357, 219–240 (1984).

13. Wilson, D.E., Scholl, B., & Fitzpatrick, D. Differential tuning of excitation and inhibition shapes direction selectivity in ferret visual cortex. Nature 560, 97–101 (2018).

14. Lien, A.D. & Scanziani, M. Cortical direction selectivity emerges at convergence of thalamic synapses. Nature 558, 80–86 (2018).

15. Billeh, Y.N., Cai, B., Gratiy, S.L., Dai, K., Iyer, R., Gouwens, N.W., Abbasi-Asl, R., Jia, X., Siegle, J.H., Olsen, S.R., Koch, C., Mihalas, S., & Arkhipov, A. Systematic Integration of Structural and Functional Data into Multi-scale Models of Mouse Primary Visual Cortex. Neuron 10, 388–403 (2020).

16. Freeman, A.W. A Model for the Origin of Motion Direction Selectivity in Visual Cortex. J. Neurosci. 41(1), 89–102 (2021).

17. Reichardt, W. Autocorrelation, a principle for the evaluation of sensory information by the central nervous system, in Sensory Communication (ed. Rosenblith, W.A.) (Wiley, New York, 1961).

18. Adelson, E.H. & Bergen, J.R. Spatiotemporal energy models for the perception of motion. J. Opt. Soc. Amer. A. 2, 284–99 (1985).

19. Emerson, R. C., Bergen, J.R. & . & Adelson, E.H. Directionally Selective Complex Cells and the Computation of Motion Energy in Cat Visual Cortex J. Neurophysiol. 32, 203–218, (1992)

20. Watson, A.B. & Ahumada, A.J. A look at motion in the frequency domain, NASA Tech. Mem. 84352 (1983).

21. Watson, A.B. & Ahumada, A.J. Model of human visual-motion sensing. J. Opt. Soc. Amer. A. 2, 322–341 (1985).

22. Hubel, D.H. & Wiesel, T.N. Receptive fields and functional architecture of monkey striate cortex. J. Physiol. 195(1), 215–43 (1968).

23. Reid, R.C. & Alonso, J.M. Specificity of monosynaptic connections from thalamus to visual cortex Nature 378, 281–4 (1995).

24. Connolly, M. & Van Essen, D. The representation of the visual field in parvicellular and magnocellular layers of the lateral geniculate nucleus in the macaque monkey. J. Comp. Neurol. 226, 544–64 (1984).

25. Silveira, L.C. & Perry, V.H. The topography of magnocellular projecting ganglion cells (M-ganglion cells) in the primate retina. Neurosci. 40, 217–37 (1991).

26. Chariker, L., Shapley. R., and Young L.S. (2016) Orientation Selectivity from Very Sparse LGN Inputs in a Comprehensive Model of Macaque V1 Cortex. J. Neurosci. 36, 12368–12384

27. Garcia-Marin, V., Kelly, J.G. and Hawken, M. J. (2019) Major Feedforward Thalamic Input Into Layer 4C of Primary Visual Cortex in Primate. Cereb. Cortex 29, 134–149 (2019)

28. Hawken, M.J., Shapley, R.M., & Grosof, D.H. Temporal frequency selectivity in monkey visual cortex. Visual Neurosci. 13, 477–492 (1996).

29. Masland, R.H. The neuronal organization of the retina. Neuron 76, 266–80 (2012).

30. Reid, R.C. & Shapley, R.M. Space and time maps of cone photoreceptor signals in macaque lateral geniculate nucleus. J Neurosci. 22, 6158–6175 (2002).

31. Jin, J., Wang, Y., Lashgari, R., Swadlow, H.A., & Alonso, J.M. Faster thalamocortical processing for dark than light visual targets. J Neurosci. 31, 17471–9 (2011).

32. DeValois, R.L. & DeValois, K.K. Spatial Vision (Oxford Univ Press, New York, 1988).

33. van Santen, J.P.H. & Sperling, G. Temporal covariance model of human motion perception. J. Opt. Soc. Am. A. 1, 451–473 (1984).

34. Tanaka, K. Cross-correlation analysis of geniculostriate neuronal relationships in cats. J Neurophysiol. 49, 1303–1318 (1983).

35. DeAngelis, G.C., Ohzawa, I., & Freeman, R.D. Receptive-field dynamics in the central visual pathways. Trends Neurosci. 18, 451–458 (1995).

36. Yeh, C.I., Xing, D., Williams, P.E., & Shapley, R.M. Stimulus ensemble and cortical layer determine V1 spatial receptive fields. Proc. Natl. Acad. Sci. U.S.A. 106, 14652–7 (2009).

37. Angelucci, A. and Sainsbury, K. (2006) Contribution of feedforward thalamic afferents and corticogeniculate feedback to the spatial summation area of macaque V1 and LGN. J. Comp. Neurol. 498, 330–51

38. Chariker, L., Shapley. R., Hawken, M., and Young L.S., in preparation (2021)

39. McLean, J. & Palmer, L.A. Contribution of linear spatiotemporal receptive field structure to velocity selectivity of simple cells in area 17 of cat. Vision Res. 29, 675–679 (1989).

40. Priebe, N.J. & Ferster, D. Direction selectivity of excitation and inhibition in simple cells of the cat primary visual cortex. Neuron 45, 133–145 (2005).

41. Marr, D. and Ullman, S. Directional selectivity and its use in early visual processing. Proc. R. Soc. London Ser. B 211, 151–180 (1981).

42. Baker, P.M. & Bair, W. Inter-Neuronal Correlation Distinguishes Mechanisms of Direction Selectivity in Cortical Circuit Models. J. Neurosci. 32, 8800–8816 (2012).

43. Moore, B. D., Alitto, H.J., & Usrey, W. M. Orientation tuning, but not direction selctivity, is invariant to temporal frequency in primary visual cortex. J. Neurophysiol. 94, 1136–1345 (2005)

44. Suarez, H., Koch, C., & Douglas, R. Modeling direction selectivity of simple cells in striate visual cortex within the framework of the canonical microcircuit. J. Neurosci. 15, 6700–6719 (1995).

45. Maex, R. & Orban, G.A. Model circuit of spiking neurons generating directional selectivity in simple cells. J. Neurophysiol. 75, 1515–1545 (1996).

46. Zhu, W., Shelley, M., & Shapley, R. A neuronal network model of primary visual cortex explains spatial frequency selectivity. J. Comput. Neurosci. 26, 271–87 (2009).

47. Ringach, D., Shapley, R.M., Hawken, M.J. Orientation selectivity in macaque V1: diversity and laminar dependence. J Neurosci. 22, 5639–5651 (2002).

48. Henry, C.A., Joshi, S., Xing, D., Shapley, R.M., Hawken, M.J. Functional characterization of the extraclassical receptive field in macaque v1: contrast, orientation, and temporal dynamics. J Neurosci. 33, 6230–42 (2013).

