## Supplemental Information for "A theory of direction selectivity for Macaque primary visual cortex"

### S1 Pref and Opp firing rates aligned at peak TF and SF.

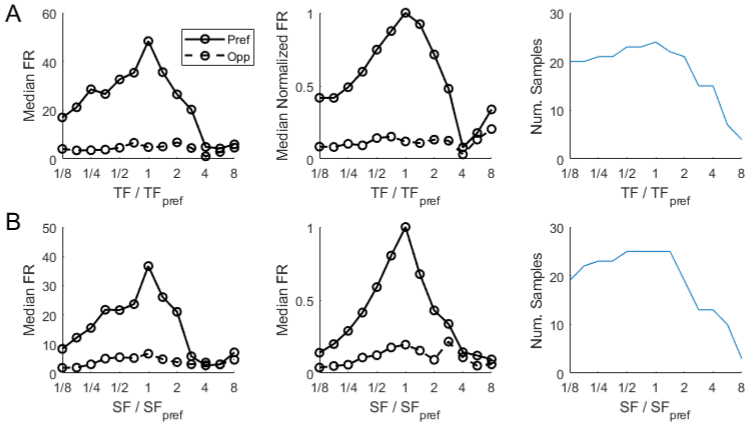


Figure S1: **Median firing rates of Pref and Opp TF and SF tuning curves aligned at peak.** Data is from the same set of cells as in figure 1 of the main text. A. Left: median Pref and Opp responses of DS $> 3$ Simple cells computed at $TF/TF_{peak}$; for each cell $TF_{peak}$ is the TF at peak in the Pref direction; Center: same as left plot, with firing rate normalized to be 1 at peak; Right: number of DS $> 3$ cells with data at each $TF/TF_{peak}$. B. Corresponding plots with SF in place of TF.

### S2 x-t plot analysis.

In figure S2, we give examples to illustrate how spatiotemporal inseparability is not a good indicator of DS for all stimuli. While a separable kernel cannot have DS, as in S2A, an inseparable one like S2B cannot guarantee DS across all temporal frequencies. There is little DS below 6 Hz in that example. Meanwhile, the inseparable kernel in S2C, with a $(2, 1)$ ON with 10 ms delay, does have DS across TF, but its x-t plot is not substantially different from that in S2B.

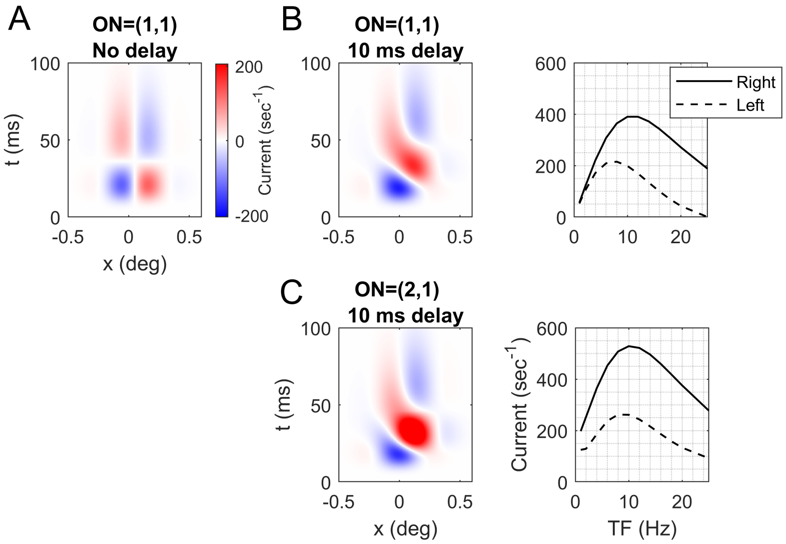


Figure S2: **x-t plots and DS response over temporal frequency.** A. Spatiotemporal kernels for pairs of OFF, ON cells separated by $d = 0.1^\circ$. In each case, the OFF cell is type (1,1). The left panel shows the separable case with ON=(1,1) with no delay; the middle panel shows ON=(1,1) with a 10 ms delay; and the righ panel shows ON=(2,1) with 10 ms delay. B. Left and Right motion responses are plotted as functions of temporal frequency, corresponding to the middle and right panels of A. SF is fixed at 2.5 c/d.
